# A spatiotemporal analysis of the effect of urbanization on birdsong

**DOI:** 10.1101/2025.10.30.685579

**Authors:** Sarah Hourihan, Nicole Creanza

**Affiliations:** Department of Biological Sciences, University of Southern California, Los Angeles, CA 90007; Department of Biological Sciences, Vanderbilt University, Nashville, TN 37212; Evolutionary Studies, Vanderbilt University, VU Box #34-1634, Nashville, TN 37235

## Abstract

Rising population density and human activities have caused an increase in low-frequency ambient noise, which disrupts auditory communication systems, such as birdsong. In oscine songbirds, birdsong is a learned behavior that functions in territory defense, species recognition, and mate attraction. Increased noise levels in urban areas can reduce signal efficacy, which can have negative impacts on fitness. However, many species have persisted in urban settings by singing at less noisy times of day, singing at higher amplitudes, or changing aspects of their song to reduce masking by low-frequency ambient noise. Previous studies have often focused on modern populations of single species in rural and urban settings; here, we conduct a spatiotemporal analysis of the effects of urbanization on the birdsongs of nine species across ∼60 years. We use machine learning to assess distinguishability between songs from historic and modern timepoints. Additionally, we assess whether variation within modern songs is associated with urbanization, and whether song differences over time are associated with changes in urbanization levels. We find that while there are some song-feature differences between historic and modern songs, and between modern songs along a rural-urban gradient, these differences are not consistent across species. These results reinforce the notion that species respond differently to urbanization by altering different aspects of their songs. For the vocal-learning songbird species studied, historic and modern songs were not distinguishable using dimensionality reduction techniques and machine learning, and some modern songs differed in more urban settings. However, the non-vocal-learning (suboscine) species showed an overall increase in its lower song frequencies in modern songs, but showed no systematic differences in song features along an urbanization gradient, suggesting more dynamic changes in the vocal learners and population-level changes in the non-learners. This spatiotemporal study is the first to compile and analyze historic and modern songs from multiple species alongside urbanization data, bringing new insights to the effects of urbanization on birdsong.

## INTRODUCTION

As human populations have grown and spread, the dramatic rise in human activities has led to modified habitats, with global ecological and evolutionary effects on other organisms [1–5]. Human density continues to increase in cities, with urban areas also expanding geographically [6–8]. As a result, low-frequency ambient noise (e.g., from vehicular traffic, aircraft, and HVAC systems) has increased to unprecedented levels [9]. Low-frequency ambient noise can mask the frequencies found in animal vocalizations, disrupting communication in songbirds (Order: Passeriformes) and other species that produce vocalizations [9,10]. In oscine birds, song is a learned behavior that functions in species recognition, mate attraction, and territory defense [11–13]. These functions can be hampered when rising noise levels reduce signal efficacy [14–18], which can result in negative fitness consequences [19]. Several studies have shown that birds experience not only lower abundance [20–23] but also reduced reproductive success in urban environments [24,25]. The patterns of reduced fitness in noisy habitats can be further compounded by difficulties in signal transmission, particularly if females prefer low-frequency songs [26,27]. Conversely, species that can tolerate a human-modified habitat can potentially have improved reproductive success because of reduced predation [28].

Many songbird species persist in urban settings [29], which prompts the question: how are these populations adapting to their rapidly changing environments? Previous focal studies have investigated how vocal signaling behaviors have adjusted to noise pollution [30,31]. The studied bird populations did not adapt their behaviors to noise in the same way; their responses include shifting to sing at less noisy times of day [32], singing at higher amplitude (i.e., the Lombard effect) [33,34], moving to a new location [35–37], and altering songs to reduce masking [30]. One of the most common song-feature changes observed in multiple species is an increase in syllable lower frequency, which leads to better transmission over low-frequency ambient noise [38–42]. However, not all avian species raise the lower frequency of their vocalizations in urban environments [42,43]. Body size and peak frequency of song are correlated [44], so some species naturally sing at higher frequencies and might not adjust their vocalizations as much in urban areas [45].

Previous studies of urban singing behaviors have mainly focused on a single species and compared songs from urban and rural populations [46] or experimentally increased noise levels to measure real-time shifts in song [47,48]. Less common are studies that include multiple species, which generally do not compare songs over a long temporal scale that would allow for study of changes in song over time and space. One study of six species across an urbanization gradient showed that two of the species significantly increased the lower syllable frequency of their songs [49]. Another study comparing the dominant frequency of song in urban and rural settings in five species demonstrated that species with intermediate dominant frequencies modified their vocalizations more than those with higher or lower dominant frequencies [42]. Finally, a study with fourteen oscine and seven suboscine species suggests that oscine species increase the lower frequency of their songs more dramatically in response to noise than suboscine species [50].

By integrating historical recordings into multi-species studies, researchers can better understand how songs are changing across several generations and over varying levels of urbanization. A multi-species temporal analysis over the scale of several decades requires an extensive database of recordings, such as the community-science repository, Macaulay Library, which was established in 1957 to catalog the growing number of recordings made possible by portable recording devices [51].

Our study is unique in exploring the evolutionary impacts of urbanization on birdsong over both time and space as well as across multiple species. We include the nine songbird species with the most historic recordings on the Macaulay Library song recording archive. We focus on the northeastern and eastern midwest regions of the United States and the southeastern region of Canada, where recordings are most dense and the species’ geographic ranges overlap (**Fig. 1**). These nine species’ songs have not been well-studied in the context of urbanization. If songs of a species have changed substantially in response to urbanization, these song differences could interfere with species recognition and mate attraction, particularly if song production has changed more quickly than song perception or preferences.

**Figure 1.**
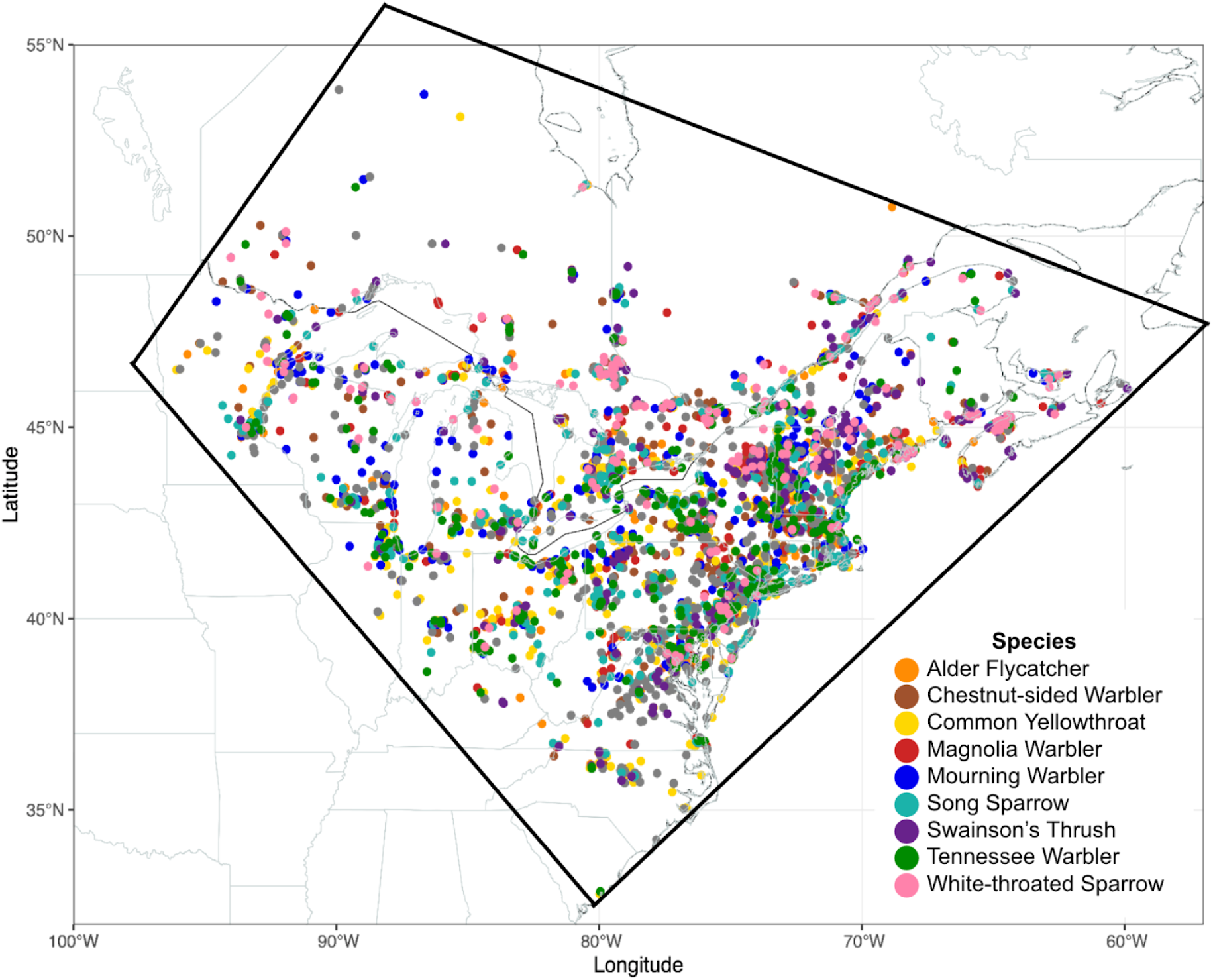
Mapped locations of the recordings included in our study. The polygon outlined is the focal area of our study. This map was created with Natural Earth in R: https://www.naturalearthdata.com/about/terms-of-use/.

Here, we explore variation between historic (from ∼1951–1963) and modern (from 2018–2023) songs of nine species. Using community-science recordings, we parsed and analyzed 2,676 songs in Chipper, a semi-automated song segmentation and analysis software [52]. Then, we extracted and analyzed song-feature data to assess how songs have changed over the course of several decades. In addition, we used QGIS to quantify urbanization metrics related to both anthropogenic light and noise, since light has also been linked to song disruptions [53,54], to assess whether song differences are associated with indicators of urbanization. We built a machine learning classifier to determine whether historic and modern songs of each species are readily distinguishable. Previously, we have investigated birdsong evolution in a single species using machine learning [55] (**Appendix 1**), but this is the first study to explore effects of urbanization on birdsong using these methods. Finally, we used generalized regression models to assess whether differences between modern and historic songs and differences in modern songs along an urbanization gradient can be explained by urbanization factors.

## METHODS

### Determining species and area to study

For this study, we aimed to determine how birdsong has changed over time in multiple species. We focused on two time ranges spaced as far apart as possible given the available data. Historic songs were recorded between ∼1951–1963 and modern songs were recorded between 2018-2023. The main criteria for species selection was sufficient number of song recordings available from the historic and modern timepoints. Digital repositories like Macaulay Library have many more songs available for recent years. Therefore, our study was restricted by the number of historic recordings available for each species. To select songbird species for the study, we downloaded all available Macaulay Library song recordings from 1963 or before and filtered results to species with ≥30 recordings. Then, we plotted the locations of those recordings on a map (**Fig. S1**). To focus on a specific area, we created a polygon around the geographical area with the most recordings available (**Fig. 1**). Then, we requested song recordings from Macaulay Library for species that had ≥20 historic song recordings taken within the polygon. After this filtering process, we identified nine species to include in the study: alder flycatcher (*Empidonax alnorum*), chestnut-sided warbler (*Setophaga pensylvanica*), common yellowthroat (*Geothlypis trichas*), magnolia warbler (*Setophaga magnolia*), mourning warbler (*Geothlypis philadelphia*), song sparrow (*Melospiza melodia*), Swainson’s thrush (*Catharus ustulatus*), Tennessee warbler (*Leiothlypis peregrina*), and white-throated sparrow (*Zonotrichia albicollis*).

### Obtaining songs for analysis

We acquired song recordings from Macaulay Library [51] by requesting all available song files for the nine species recorded within the historic and modern periods. For our analyses, we compiled metadata for each recording, including species, latitude, longitude, date, and time. These details are generally provided by the recordist when uploading recordings to Macaulay Library. For recordings that specified a location but no geographic coordinates, we approximated the latitude and longitude by searching the location in Google Maps.

We used Audacity (version 3.1.3) to extract one song from each recording at a sampling rate of 44,100 Hz (**Fig. 2A**). Recordings with no suitable song bouts due to background noise were omitted from the study.

**Figure 2.**
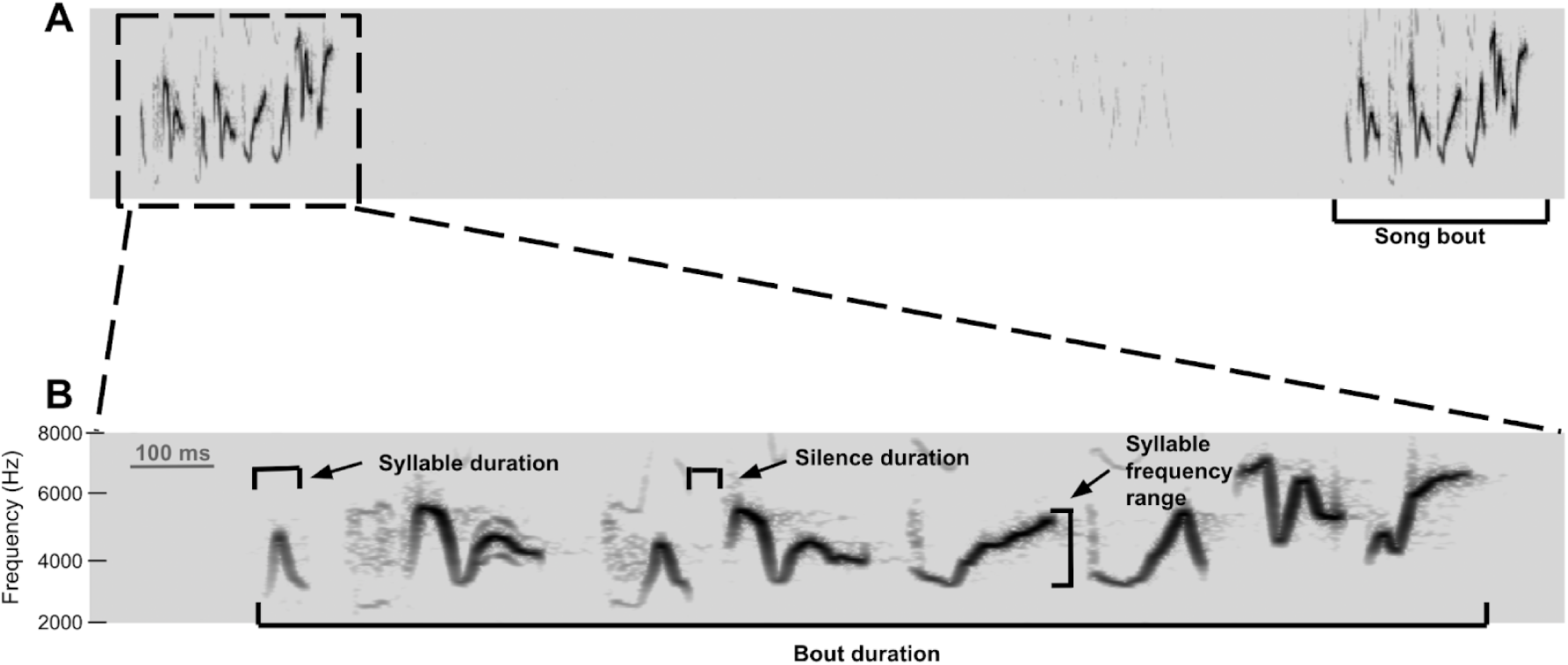
(A) Example spectrogram of a song recording from Macaulay Library with multiple song bouts (magnolia warbler, ID Number: 9727). (B) Example spectrogram of an extracted song. Songs can be segmented into syllables and are characterized by song features, such as bout duration, average syllable and silence duration, and average syllable frequency range.

### Song segmentation and quantification of song features

To parse and analyze songs, we used Chipper [52], a graphical user interface that first allows users to supervise an automated syllable segmentation process and then analyzes properties of those syllables. Chipper parses songs into syllables by identifying periods of sound separated by silence. To prevent lower amplitude syllables from being segmented incorrectly, we normalized amplitude across each song. To improve segmentation, we adjusted the signal-to-noise threshold, minimum syllable duration, and minimum silence duration. If any syllables were incorrectly parsed after these adjustments, we manually modified the syllable segmentation. If a song could not be cleanly segmented due to poor audio quality or excessive background noise, the song was excluded from the study.

After song segmentation, we examined a random subset of 20 songs from each species—10 historic songs and 10 modern songs—to estimate the noise threshold, defined as the minimum size that a cluster of sound must be in order to not be discarded as noise [56]. This threshold primarily affects syllable-level song features, such as average syllable frequency range, average syllable lower frequency, and average syllable upper frequency. Additionally, we used the same subset of songs to estimate the syllable similarity threshold, defined as the percent syllable overlap that determines whether two syllables are considered the same or different [56]. This threshold primarily affects song features related to syntax, such as the number of unique syllables. The estimated noise and syllable similarity thresholds for each species are in **Table S1**. Species’ estimated noise and syllable similarity thresholds were subsequently applied for analysis of all songs of each respective species.

Lastly, we used the song analysis function in Chipper, which considers the syllable segmentation information and the noise and syllable similarity thresholds to generate quantitative song-feature data. We exported nine song-feature outputs from Chipper for our analyses: average syllable duration (measured in milliseconds, ms), bout duration (ms), number of syllables, rate of syllable production (1/ms, measured as number of syllables divided by bout duration), average silence duration (ms), number of unique syllables, average syllable upper frequency (measured in Hertz, Hz), average syllable lower frequency (Hz), and average syllable frequency range (Hz).

### Analyses comparing historic and modern song-feature data

To reduce dimensionality of song-feature data, we used principal component analysis (PCA), linear discriminant analysis (LDA), uniform manifold approximation and projection (UMAP), and t-distributed stochastic neighbor embedding (t-SNE) on the song-feature data of each species. These analyses visualize song-feature variation across the nine studied song features and show whether song data clusters by timepoint (i.e., ‘historic’ or ‘modern’) in two-dimensional space.

We fit generalized linear models (GLMs) to determine whether ‘historic’ and ‘modern’ labels are informative of song-feature variation. To estimate the amount of urbanization that has occurred between the historic and modern timepoints, we determined percent population density change using U.S. and Canadian census data. We quantified the percent population change in each state or province from 1960 to 2020 in the U.S and from 1961 to 2021 in Canada [57,58]. Additionally, we estimated the change in nighttime light intensity using NOAA’s composite nighttime light images with 30 arc-second grids from the earliest and latest comparable maps (1992 and 2013). To extract nighttime light intensity for the specific locations of our song recordings, we used QGIS [59]. We subtracted 1992 nighttime light intensity from 2013 nighttime light intensity to calculate the change in nighttime light intensity at each recording’s location.

We fit GLMs separately for each species and each of their nine studied song features. Each model included timepoint category (historic or modern), change in population density, and change in nighttime light intensity as main effects. Interaction terms between timepoint category and each of the two continuous urbanization metrics were included to test whether the effects of change in population density and change in nighttime light intensity differ for modern songs with historic songs as the reference group. The model summaries were used to identify which effects are significantly associated with variation in a given song feature within a given species.

### Machine learning classifier to distinguish historic and modern songs

We built a random forest machine-learning classifier using the tidymodels meta-package in R to assess how accurately historic and modern songs of each species can be distinguished [60]. For each species, 75% of the songs were randomly selected to train the model and the remaining 25% of songs were used to test the model’s accuracy. We downsampled the training set using the themis R package [61] to obtain a balanced sample of songs from the historical and modern time ranges. The classifier uses the nine studied song features and 100 decision trees to make predictions of ‘historic’ or ‘modern’ for test song classification. When we increased the number of decision trees, we found that the classifier’s prediction accuracy was largely unaffected (**Table S2**). We tuned the hyperparameters ‘mtry’ (number of predictors that are randomly sampled at each split when creating the tree models) and ‘min_n’ (minimum number of data points in a node that are required for the node to be split further) for each round of classification. For each species, we used the vip R package [62] to determine which song features were most valuable for making the predictions.

### Obtaining urbanization metrics data for the analysis of modern songs

For the analysis of modern songs along an urbanization gradient, we quantified five urbanization metrics at each modern song recording’s location: population density, nighttime light intensity, tree coverage, distance to the nearest major road, and a composite metric called the Human Footprint Index.

For population density, we used the Gridded Populations of the World map (version 4) from NASA’s Socioeconomic Data and Applications Center [63]. The map contains estimates of human population density (persons per km^2^) in 2015 based on national censuses and population registries with respect to relative spatial distribution at 30 arc-second resolution.

For nighttime light intensity, we used the Nighttime Lights Time Series from NOAA’s National Centers for Environmental Information. The composite nighttime light images with 30 arc-second grids were captured by visible and infrared light sensors flown on Defense Meteorological Satellite Program satellites in 2013.

For tree coverage, we used the Global Forest Change map used by Hansen et al. [64], which contains information on the percentage of canopy closure in 2000, such that trees are defined as vegetation taller than 5 meters in height, pixels represent an approximately 30 meter grid, and a pixel value of 0 indicates no tree cover and 100 indicates full tree cover.

For the Human Footprint Index, we extracted values from the 2009 Human Footprint map from Venter et al. [65,66][66] which estimated the global human influence on the environment as a weighted composite of multiple factors: (1) extent of built environments; (2) crop land; (3) pasture land; (4) human population density; (5) night-time lights; (6) railways; (7) roads; and (8) navigable waterways. These metrics were weighted according to estimates of their levels of human pressure and then summed to create composite values.

For distance to the nearest major road, we used the Roads map (version 5.0.0) from Natural Earth, which displays roads from CEC North America Environmental Atlas at a 10 meter scale. Major roads include major and secondary highways, beltways, and ferry routes.

To extract urbanization data for the specific locations of our song recordings, we used QGIS. First, we added the raster map layers of our urbanization metrics. Then, we added a delimited text layer with the latitude-longitude coordinates of our song recordings. For population density, nighttime light intensity, tree coverage, and Human Footprint Index, we used the “Sample raster values” tool to obtain data for those metrics at each song recording’s coordinates. For distance to the nearest major road, we used the “Join attributes by nearest” tool to obtain the distance between each recording’s coordinates and the nearest major road.

### Assessing the relationship between urbanization and modern song-feature data

First we tested the correlation between the urbanization metrics with pairwise correlation tests to ensure that the metrics capture different aspects of urbanization. Nighttime light intensity and Human Footprint Index were strongly correlated for the locations of our recordings (*r* = 0.89), so we only included nighttime light intensity in our models. To understand the relationship between urbanization and modern-day song-feature variation, we used generalized linear mixed models (GLMMs) fit by maximum likelihood. We fit models separately for each species and each of their nine studied song features using tree coverage, population density, distance to the nearest major road, and nighttime light intensity as fixed effects and month of recording as a random effect. We set the GLMM family as Gaussian, with a log link function. To ensure model convergence, we used the “bobyqa” optimizer and set the maximum number of function evaluations to 2^8^. The model summaries were used to identify which fixed effects are significantly associated with variation in a given song feature within a given species.

## RESULTS

### Determining species and area to study

The focal area of our study was the northeastern and eastern midwest regions of the United States and the southeastern region of Canada (**Fig. 1**). This area has the most historic song recordings available and contains locations with varying levels of urbanization, making it an ideal area to study. In our study, we included the nine species with the most historic recordings available within the focal area: alder flycatcher (*Empidonax alnorum*), chestnut-sided warbler (*Setophaga pensylvanica*), common yellowthroat (*Geothlypis trichas*), magnolia warbler (*Setophaga magnolia*), mourning warbler (*Geothlypis philadelphia*), song sparrow (*Melospiza melodia*), Swainson’s thrush (*Catharus ustulatus*), Tennessee warbler (*Leiothylpis peregrina*), and white-throated sparrow (*Zonotrichia albicollis*). Although there are fewer historic recordings for each species, the recording locations of the historic and modern songs of each species are generally distributed throughout the polygon (**Fig. S2**), allowing for comparison of historic and modern songs. The alder flycatcher is a suboscine species, while the rest are oscine species. Each of the species has a different species-typical song (**Fig. 3**).

**Figure 3.**
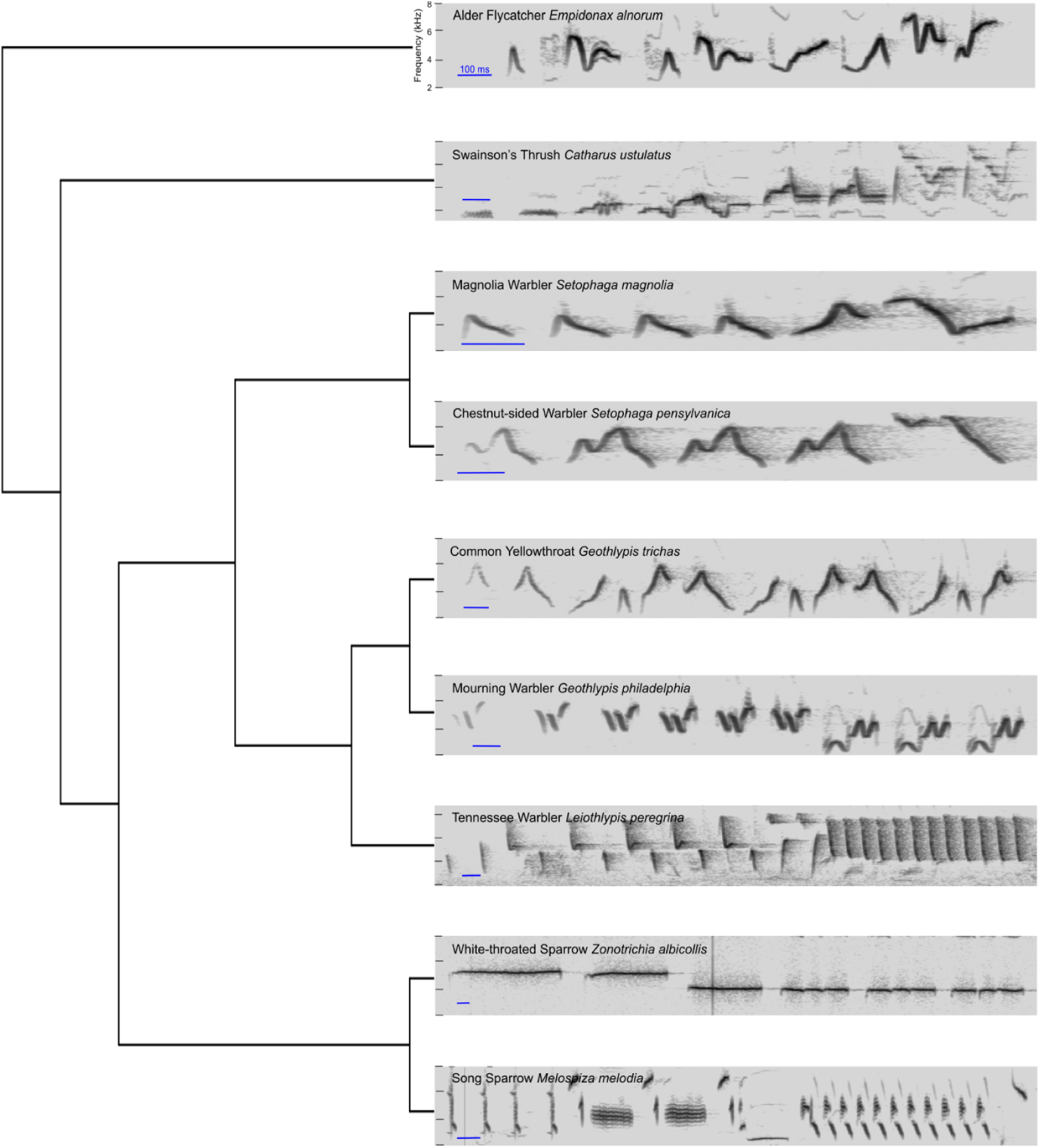
Phylogenetic tree of the nine species included in this study. At each branch tip is a representative spectrogram of the species’ song.

### Obtaining songs for analysis

In total, we obtained 4,746 song recordings from Macaulay Library. Recordings with poor audio quality or excessive background noise were excluded. We extracted one song from each of the usable recordings, resulting in 2,676 songs included in the study (**Table 1**).

**Table 1.**
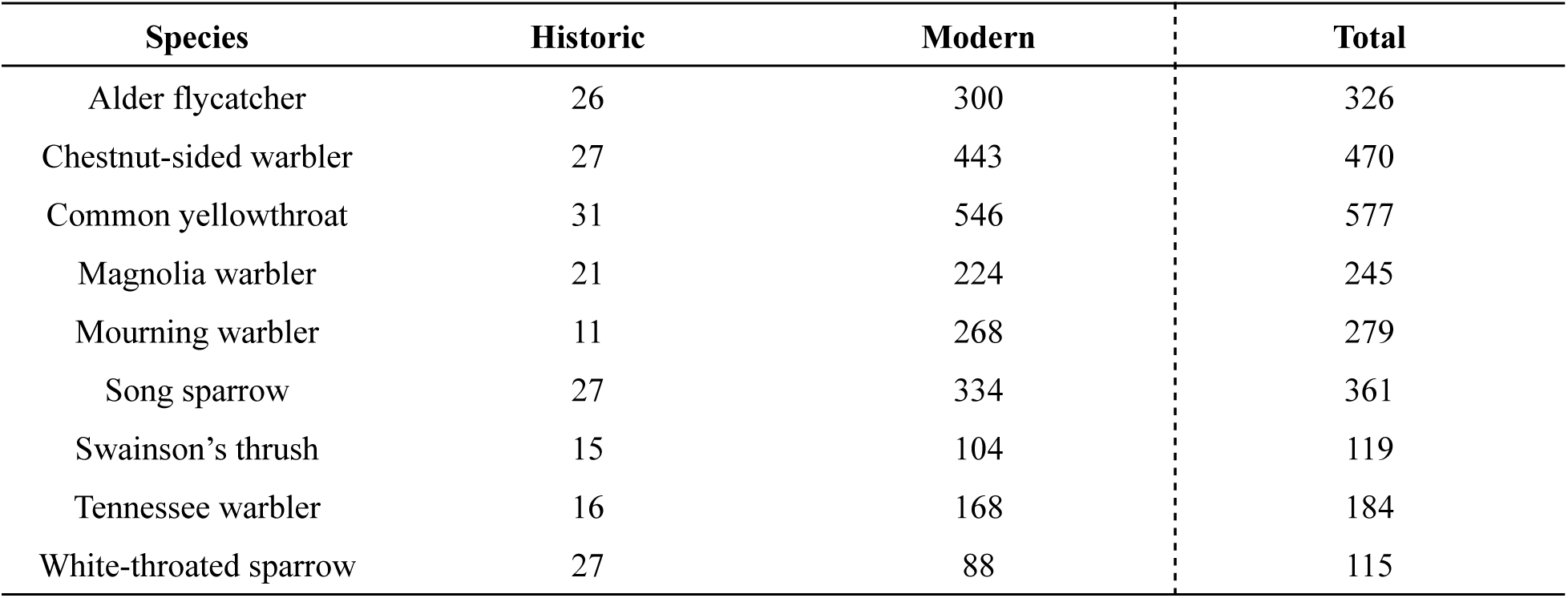
The number of historic (recorded ∼1951–1963) and modern (recorded 2018–2023) songs per species included in the study.

### Analysis of quantitative song-feature data

After segmenting and analyzing songs in Chipper, we generated data for nine song features: average syllable duration (ms), bout duration (ms), number of syllables, rate of syllable production (1/ms), average silence duration (ms), number of unique syllables, average syllable upper frequency (Hz), average syllable lower frequency (Hz), and average syllable frequency range (Hz).

Principal component analyses (PCA) of the song-feature data revealed considerable overlap in song variation among historic and modern songs of all nine species (**Fig. 4**). The proportion of each species’ song-feature variance that can be explained by principal component 1 (PC1) and principal component 2 (PC2) ranged from 33.09–45.25% and 17.27–27.20%, respectively (**Table S3**). Visualization of each species’ song-feature data with linear discriminant analysis (LDA), uniform manifold approximation and projection (UMAP), and t-distributed stochastic neighbor embedding (t-SNE) showed similar patterns, with no distinct clusters for historic and modern songs (**Fig. S3-S5**). Lack of clusters using multiple methods of dimensionality reduction suggests that the historic and modern songs of each species share a considerable amount of song-feature variation and might not be readily distinguishable.

**Figure 4.**
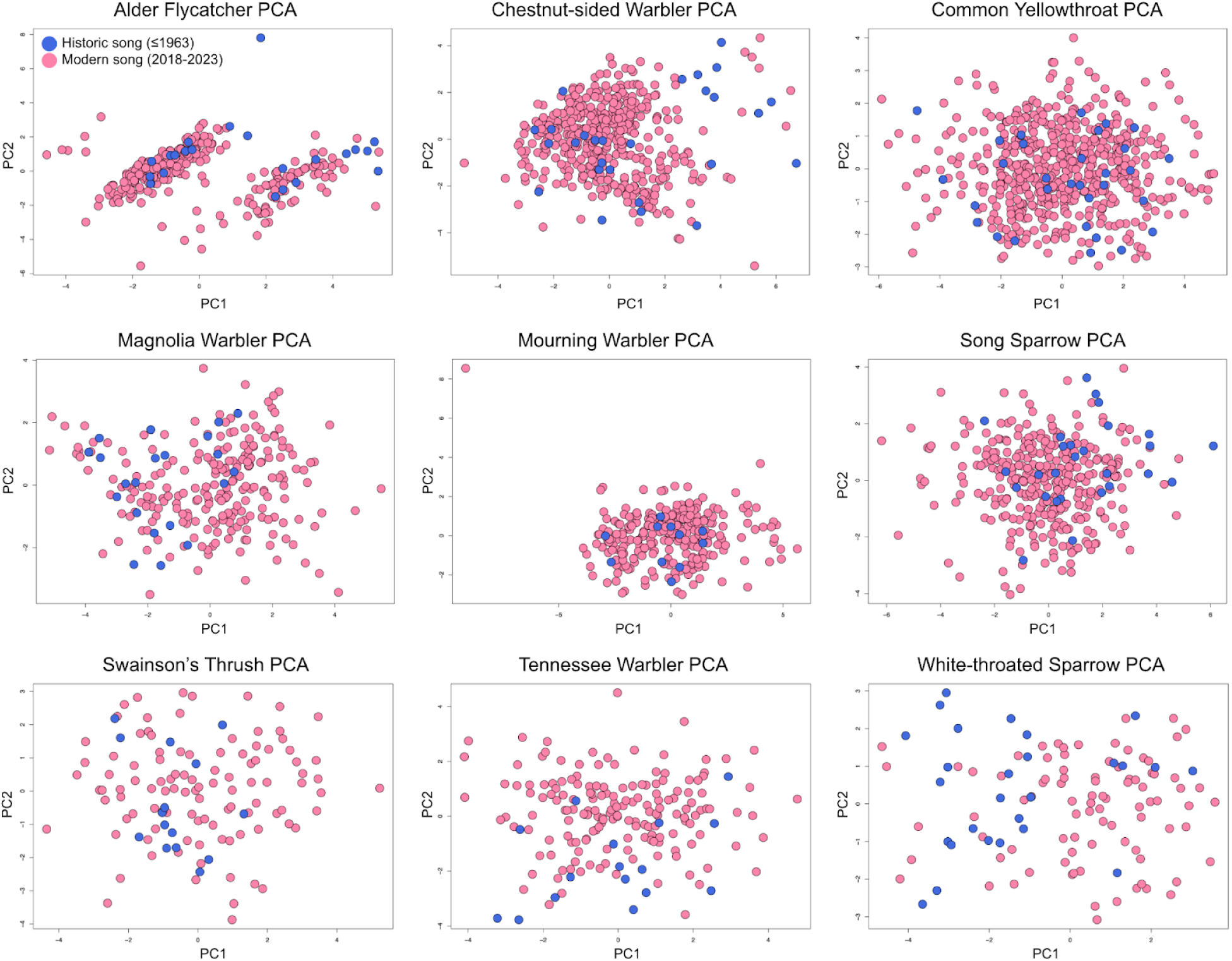
Visualization of PC1 and PC2 of each species’ song-feature data. No distinct clusters formed in PC space for the historic and modern songs.

Using generalized linear models (GLMs), we determined that ‘historic’ and ‘modern’ categorization is informative for at least one of the studied species in all nine features (**Fig. 5A**). However, the results are inconsistent across species and song features. The strongest finding is within song sparrows, with time classification associated with variation in five of the nine studied song features. Within song sparrows, time classification was informative for both syllable- and song-level features. Change in population density from 1960 to 2020 was associated with differences between modern and historic songs for specific song features within select species, particularly the magnolia warbler (**Fig. 5B**). This urbanization metric did not have consistent predictive capability for any particular species or song feature. Change in nighttime light intensity was only associated with variation between the number of syllables in modern and historic songs of the song sparrow (**Fig. 5C**). Details of GLM results and the directions of significant effects can be found in **Table S4.**

**Figure 5.**
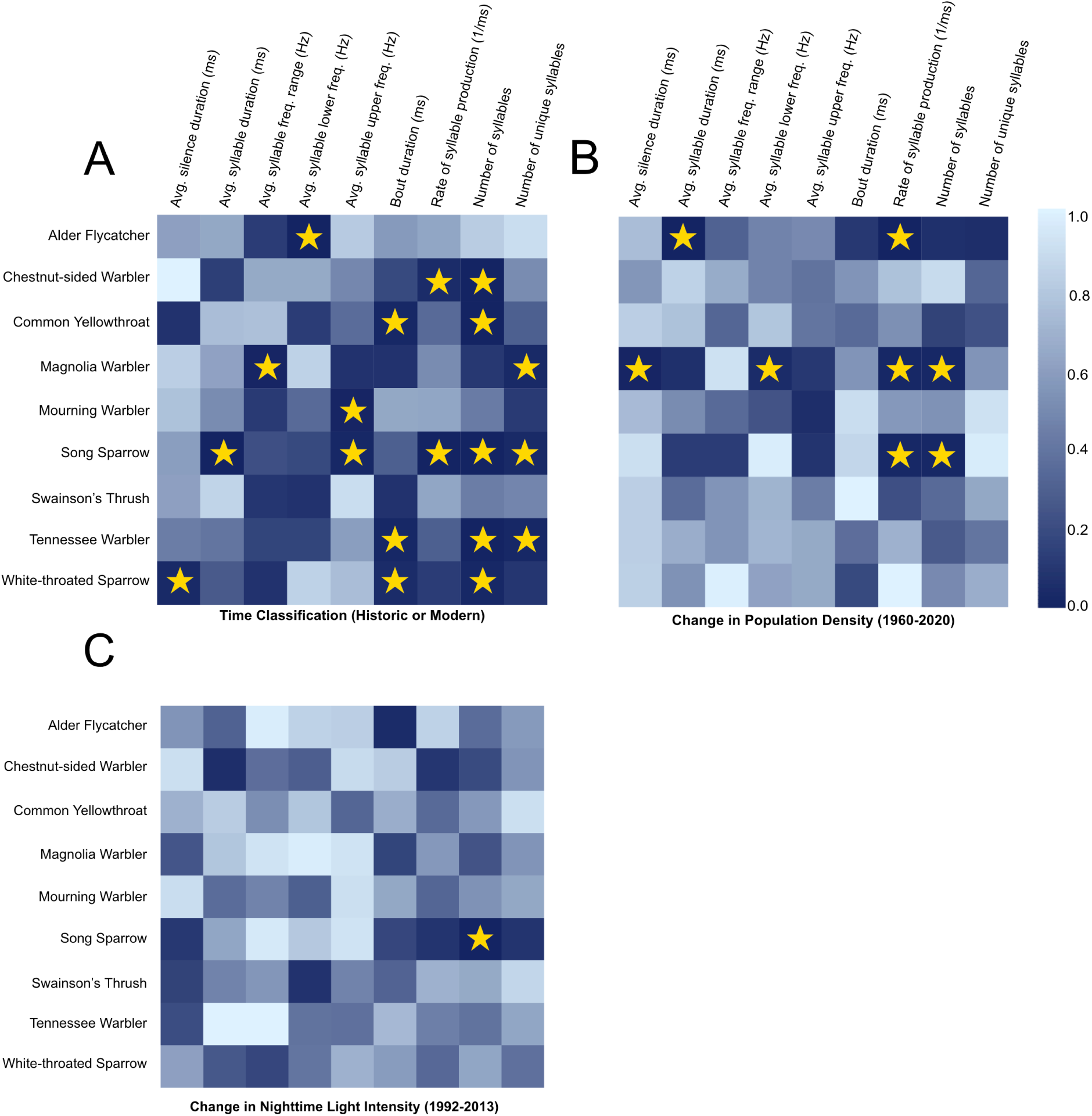
Results from the GLMs displayed as heat maps for each fixed effect. The color scale indicates *P*-value, with darker colors indicating lower *P*-values. Significant fixed effects are denoted with stars. (A) *P*-values for time classification’s effect on song-feature values across species. (B) *P*-values for change in population density’s effect on modern compared to historic song-feature values across species. (C) *P*-values for change in nighttime light intensity’s effect on modern compared to historic song-feature values across species.

### Machine learning classifier

The random forest machine-learning classifier made relatively reliable predictions of whether songs were historic or modern for some species (e.g., white-throated sparrow, 88.7%) but made predictions at or worse than random chance for others (e.g., mourning warbler, 34.6%; common yellowthroat, 50.4.%; Tennessee warbler, 50.4%) (**Table 2**). The sizes of the training sets varied due to the number of recordings available for each species. However, the classifier’s performance was not significantly correlated with training set size (**Fig. S6**). The variables of importance for model predictions differed for each species, but the most common song feature the classifier used to discriminate songs of historic or modern timepoints was average silence duration (**Table S5**).

**Table 2.** The random forest classifier’s accuracies for distinguishing historic and modern songs of each species.

| Species | Training Set Size | Historic Accuracy | Modern Accuracy | Balanced accuracy |
| --- | --- | --- | --- | --- |
| Alder flycatcher | 18 | 0.500 | 0.800 | 65.0% |
| Chestnut-sided warbler | 18 | 0.778 | 0.688 | 73.3% |
| Common yellowthroat | 23 | 0.250 | 0.757 | 50.4% |
| Magnolia warbler | 15 | 0.500 | 0.768 | 63.4% |
| Mourning warbler | 8 | 0.333 | 0.358 | 34.6% |
| Song sparrow | 19 | 0.750 | 0.542 | 64.6% |
| Swainson's thrush | 11 | 1.00 | 0.615 | 80.8% |
| Tennessee warbler | 13 | 0.333 | 0.674 | 50.4% |
| White-throated sparrow | 20 | 1.000 | 0.773 | 88.7% |

### Urbanization metrics and analysis for modern songs along an urbanization gradient

The metrics we used for analysis of modern songs along an urbanization gradient were distance to the nearest major road, nighttime light intensity, local population density, and tree coverage. None of these urbanization metrics were strongly correlated with one another (**Table S6**), suggesting that each captures different aspects of urbanization. The direction of association between the urbanization metrics reflects our expectations that nighttime light intensity and local population density are positively correlated with urbanization whereas tree coverage and distance to the nearest major road are negatively correlated with urbanization.

GLMMs fit to the modern song data revealed that certain urbanization metrics are significantly associated with variation in select song features in select species (**Fig. 6**). For example, in the common yellowthroat, songs recorded closer to a major road had shorter average silence durations and larger frequency ranges, and average syllable upper frequency increased with less tree coverage. In the white-throated sparrow, both the lower and upper frequencies of syllables increased in areas with more light pollution. Details of GLMM results and the directions of significant fixed urbanization effects can be found in **Table S7.** Visualizations of a subset of the significant GLMM results show that when an urbanization metric has a significant effect on a song feature, the given urbanization metric and song feature are weakly correlated (**Fig. 7A-C**).

**Figure 6.**
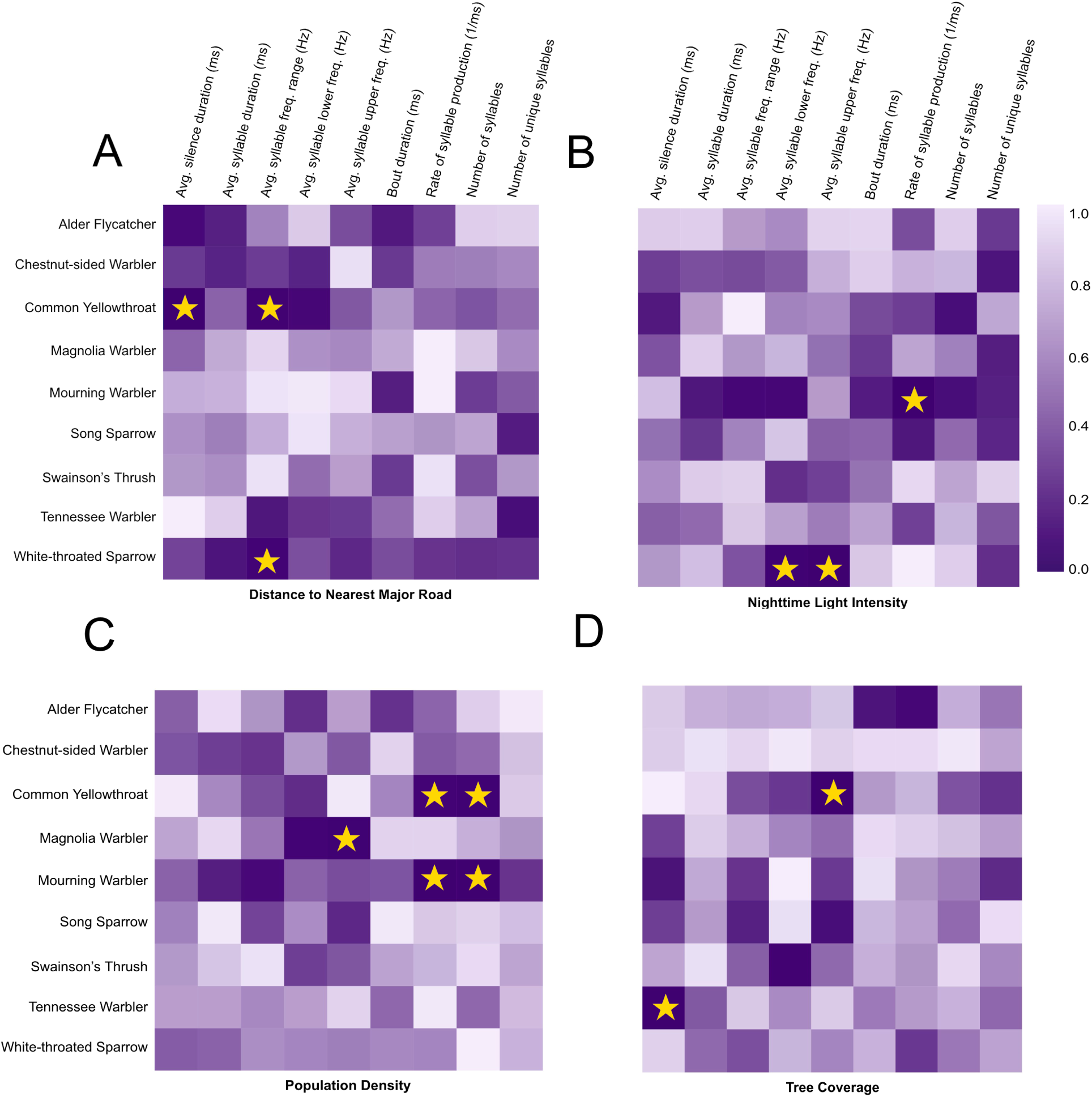
Results from the GLMMs displayed as heat maps for each urbanization metric. The color scale indicates *P*-value, with darker colors indicating lower *P*-values. Significant fixed effects are denoted with stars. (A) *P*-values for distance to the nearest major road’s effect on song-feature values across species. (B) *P*-values for nighttime light intensity’s effect on song-feature values across species. (C) *P*-values for population density’s effect on song-feature values across species. (D) *P*-values for tree coverage’s ability effect on song-feature values across species.

**Figure 7.**
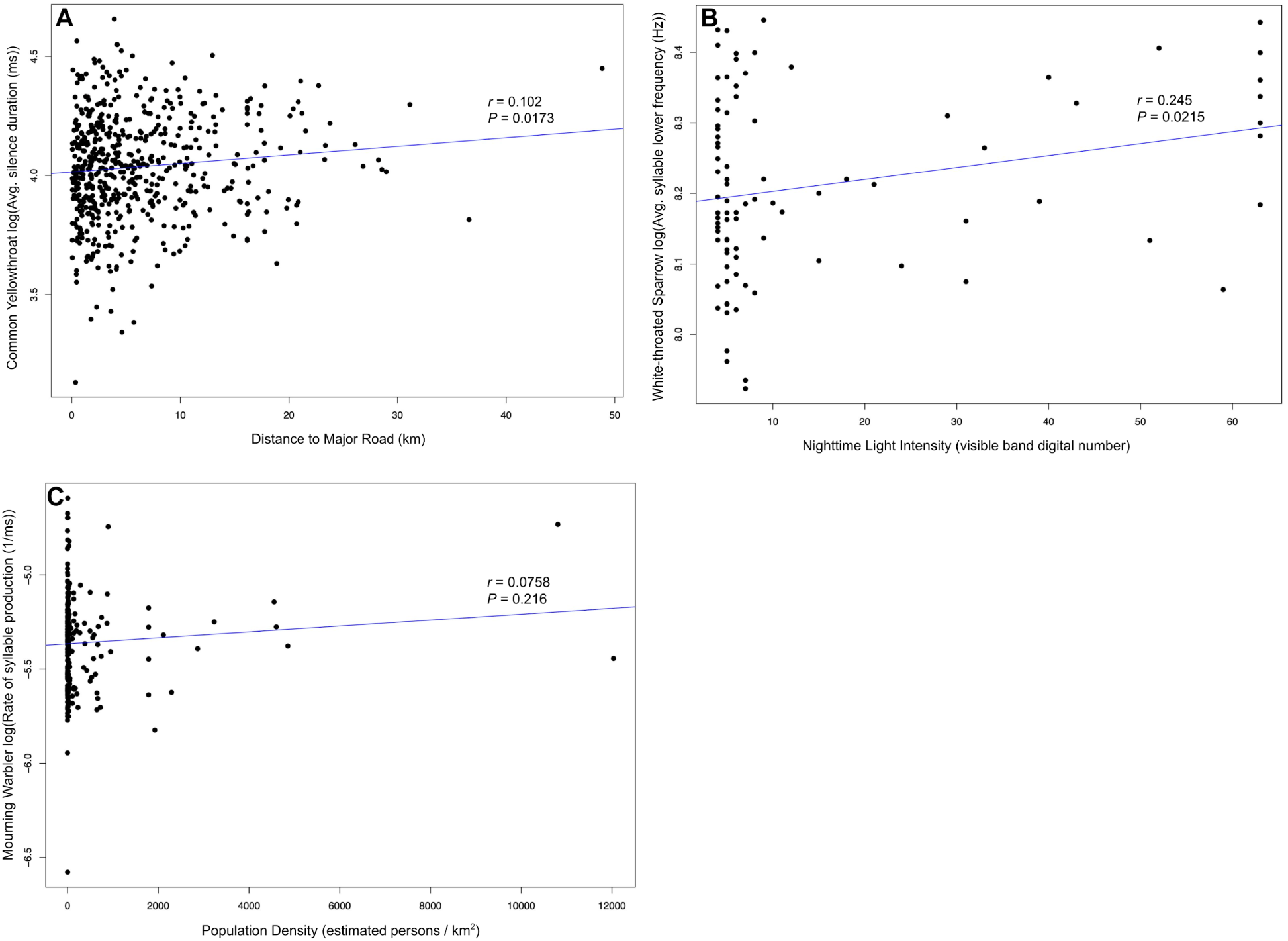
Visualizations for a subset of the significant GLMM results. (A) ‘Distance to Major Road’ had a significant effect on the common yellowthroat’s average silence duration (*P* = 0.00425); these variables have a weak positive correlation (*r* = 0.102). (B) ‘Nighttime Light Intensity’ had a significant effect on the white-throated sparrow’s average syllable lower frequency (*P* = 0.024); these variables have a weak positive correlation (*r* = 0.245). (C) ‘Population Density’ had a significant effect on the mourning warbler’s rate of syllable production (*P* = 0.00251); these variables have a weak correlation (*r* = 0.0758).

## DISCUSSION

Previous studies on the effects of urbanization on birdsong have mainly focused on songs from modern-day urban and rural areas. In urban areas, some species, including song sparrows [40], great tits [38], and house finches [67], have been shown to increase the lower frequency of their songs to improve acoustic transmission amidst high levels of low-frequency ambient noise. However, not all avian species raise the lower frequency of their vocalizations in urban environments [42,43], so this pattern cannot be generalized across the songbird clade. These prior studies show that the upper frequencies of songs are less likely to change than the lower ones. The upper frequencies of songs are likely constrained by morphological and kinematic factors, such as body size, head angle, beak shape, and beak gape [44,68–70]. Additionally, the upper frequencies of songs are not as masked by low-frequency urban noise, reducing pressure to alter them. Another adjustment observed in several avian populations living in urban areas is raising vocalization amplitude, thus increasing signal-to-noise ratio (i.e., the Lombard effect) [33,71,72].

Few studies have investigated how rising noise pollution and urbanization affect song features of multiple bird species [42,49,50] and across decades of time [73]. A majority of previous studies focus on a single species across a local urbanization gradient; these studies generally have the statistical power to detect subtle song differences in response to variation in anthropogenic noise levels, but their species-specificity presents a challenge for applying the results broadly across the songbird clade. Here, we use machine learning, community-science recordings, and statistical models to examine how birdsong differs across time, space, and species in the face of varying levels of urbanization.

All species in our study are from the oscine suborder of passerine birds, except the alder flycatcher, which belongs to the suboscine suborder. Oscine species learn their vocalizations, while suboscine species have innate songs. This confers a potential advantage to oscine populations in urban habitats, as they can adapt more quickly to their local acoustic environments. Therefore, suboscines may be more vulnerable to urban noise pollution and less capable of establishing populations in cities. However, there is some behavioral plasticity (within morphological and kinematic limits) that allows individuals to make immediate adjustments to song frequency and amplitude in response to heightened environmental noise [28,74]. The modern songs of the suboscine species in our study, the alder flycatcher, did not show any associations with the four included urbanization metrics (**Fig. 6**). However, the modern songs of the alder flycatcher showed an increase in average syllable lower frequency compared to historic songs, suggesting that the unlearned songs of this species might have changed to avoid competing with anthropogenic noise (**Fig. 5A**). In addition, the modern songs were faster in locations that had experienced greater changes in population density, showing a decrease in average syllable duration and increase in rate of syllable production compared to historic songs (**Fig. 5B**).

The dimensionality reduction analyses showed substantial overlap in song-feature variation between songs across all timepoints, with no formation of distinct historic and modern clusters in two-dimensional space for any of the dimensionality reduction methods used (**Fig. 4, Fig. S3-S5**). The GLMs revealed that time categorization (‘historic’ or ‘modern’) was informative for some species and song features, but with no consistent or generalizable patterns in either (**Fig. 5A**). The classification of historic and modern songs using machine learning mirrors these varied results, with some species having little to no differentiation between historic and modern songs and other species having stronger distinguishability. Distinguishability between historic and modern songs could have implications for mate attraction. In some species, current birds prefer modern songs over historic songs [73], but in other species, females have been shown to prefer song characteristics that have changed in response to urbanization, such as lower frequency singing [26,27], and these conflicting preferences could be a mechanism for population decline in urban areas. Perception of acoustic signals in urban environments is less studied than acoustic signaling; however, a focal study of great tits (*Parus major*) showed that receivers can adapt to detect higher frequency songs when exposed to low-frequency noise [75]. In future studies, signal perception and female choice for the nine species of this study could be evaluated in the field with playback experiments using historic and modern songs.

Using GLMMs, we evaluated the relationship between urbanization and modern song-feature data using metrics of population density, nighttime light intensity, tree coverage, and distance to the nearest major road. These metrics are proxies for the level of anthropogenic change to the environment and are meant to capture the potential disruptions birds may face due to expansion of urban areas and population growth. We included multiple measures of urbanization, because a previous study has demonstrated that species appear to have varied responses to different urbanization features [76]. Therefore, it is important to account for multiple metrics to capture a more representative range of responses. Overall, the urbanization metrics were not consistently associated with song-feature changes across the nine species (**Fig. 6**). However, these metrics were more informative for some species than others. For example, cumulatively, the urbanization metrics had significant effects on five of the common yellowthroat’s nine studied song features. In cases where an urbanization metric had a significant effect on a certain species and song feature, the correlation between the respective urbanization metric and song feature was generally weak (**Fig. 7A-C**).

This study reinforced results from previous studies that some species alter their song frequencies in response to urban noise [38–42,49,67]. We found evidence that magnolia warblers, white-throated sparrows, and common yellowthroats might sing at higher frequencies in areas that are more urban by one or more of our metrics. Interestingly, comparing modern to historic songs, we only saw an overall increase in syllable lower frequency in the alder flycatcher, the only species with unlearned song, potentially suggesting a selection pressure favoring songs that do not overlap with low-frequency noise. In contrast, in oscine species with learned song, there was no overall difference in lower frequency between modern and historic songs but several associations between increased frequency and metrics of urbanization, which could imply that oscine songbirds are more likely to dynamically increase their song frequency in the presence of low-frequency noise.

In summary, we find that although there are a handful of song-feature-level differences observed across long timescales for the nine studied species, collectively the differences are not substantial enough to reliably differentiate historic and modern songs using dimensionality reduction or machine learning classification. Our results suggest largely overlapping variation between rural and urban songs across time and space, with little consistency in how a given species’ song features will shift in response to increased urbanization. However, we did observe some song differences that are consistent with previously reported trends. Therefore, it is possible that most of the previously observed patterns of song change along an urbanization gradient, such as increased lower syllable frequency, are primarily detectable through comparisons at a local level and might not scale to patterns that are detectable at a population level or across large distances. Our findings thus suggest that the song differences that are detectable across large spatiotemporal scales could indicate which song features are broadly important for signaling in those species.

## ACKNOWLEDGEMENTS

We are grateful for song recording media from The Macaulay Library at the Cornell Lab of Ornithology (www.macaulaylibrary.org) and for the use of publicly available images processed by NOAA’s National Geophysical Data Center. Thank you to members of the Creanza Laboratory for their feedback on this project.

