## Supplemental Materials for "A spatiotemporal analysis of the effect of urbanization on birdsong"

**Supplemental Materials: Hourihan and Creanza (2025). A spatiotemporal analysis of the effect of urbanization on birdsong.**

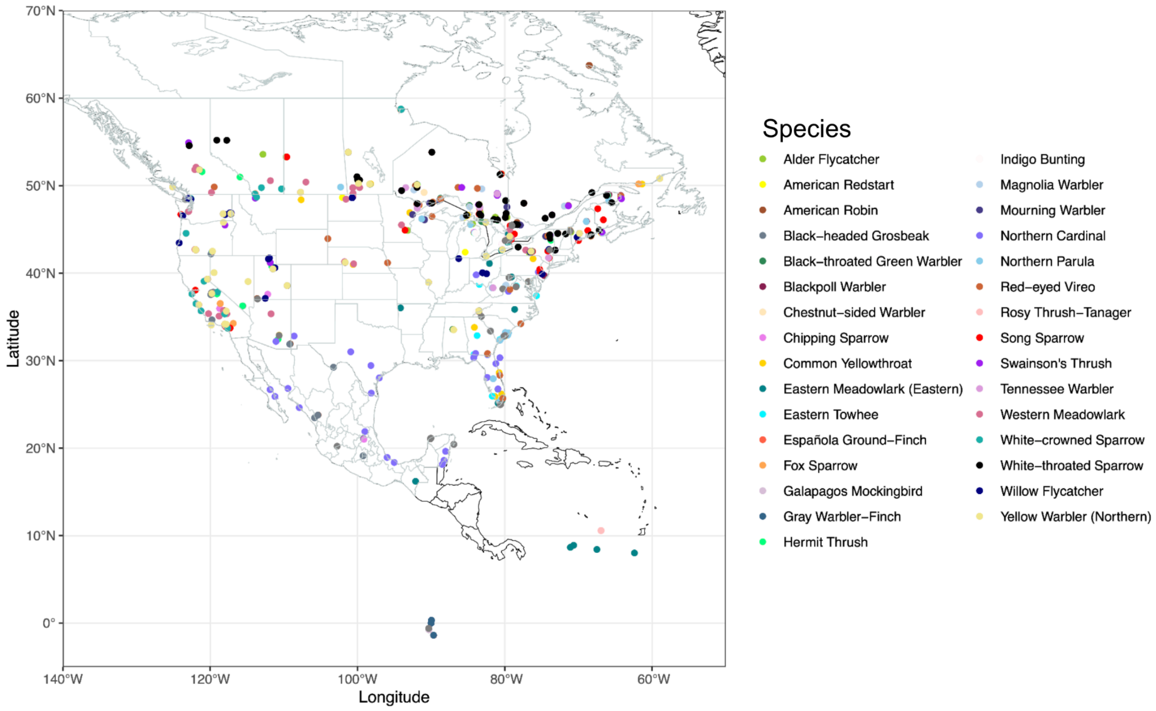


**Figure S1** The recording locations of candidate species that had ≥30 historic recordings. Using this map, we chose our focal area to be the northeastern and eastern midwest region of the United States and the southeastern region of Canada. This map was created with Natural Earth in R: https://www.naturalearthdata.com/about/terms-of-use/.

**Table S1** The noise and syllable similarity thresholds applied to each species for song analysis in Chipper.

| Species | Noise Threshold (# of Matrix Elements) | Syllable Similarity Threshold (%) |
|---|---|---|
| Alder flycatcher | 79 | 36.5 |
| Chestnut-sided warbler | 128 | 41.0 |
| Common yellowthroat | 113 | 45.8 |
| Magnolia warbler | 109 | 40.5 |
| Mourning warbler | 90 | 32.6 |
| Song sparrow | 97 | 48.5 |
| Swainson's thrush | 79 | 38.0 |
| Tennessee warbler | 115 | 45.6 |
| White-throated sparrow | 116 | 46.5 |

**Table S2** The random forest classifier's discrimination between the alder flycatcher's historic and modern songs is largely unaffected by increasing the number of decision trees. Therefore, we used 100 decision trees for our random forest classifiers.

| Number of decision trees | Historic accuracy | Modern accuracy | Balanced accuracy |
|---|---|---|---|
| 100 | 0.500 | 0.800 | 65.0% |
| 150 | 0.500 | 0.787 | 64.4% |
| 200 | 0.500 | 0.787 | 64.4% |
| 500 | 0.500 | 0.787 | 64.4% |
| 1000 | 0.500 | 0.800 | 65.0% |
| 2000 | 0.500 | 0.813 | 65.7% |

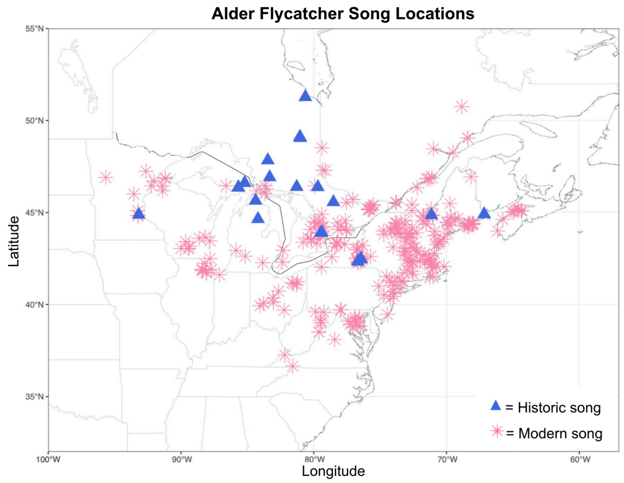
Alder Flycatcher Song Locations
55°N
50°N
45°N
40°N
35°N
Latitude
100°W
90°W
80°W
70°W
60°W
Longitude
= Historic song
= Modern song

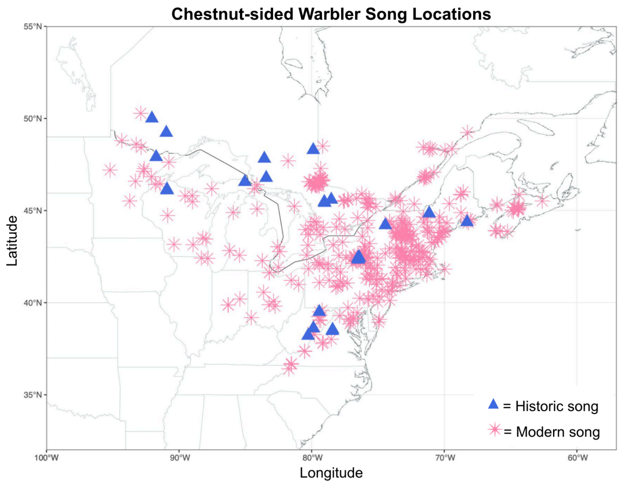
Chestnut-sided Warbler Song Locations
55°N
50°N
45°N
40°N
35°N
Latitude
100°W
90°W
80°W
70°W
60°W
Longitude
= Historic song
= Modern song

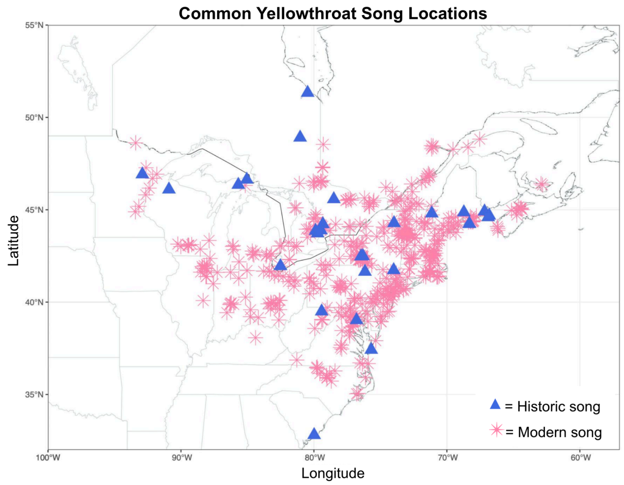
Common Yellowthroat Song Locations
55°N
50°N
45°N
40°N
35°N
Latitude
100°W
90°W
80°W
70°W
60°W
Longitude
= Historic song
= Modern song

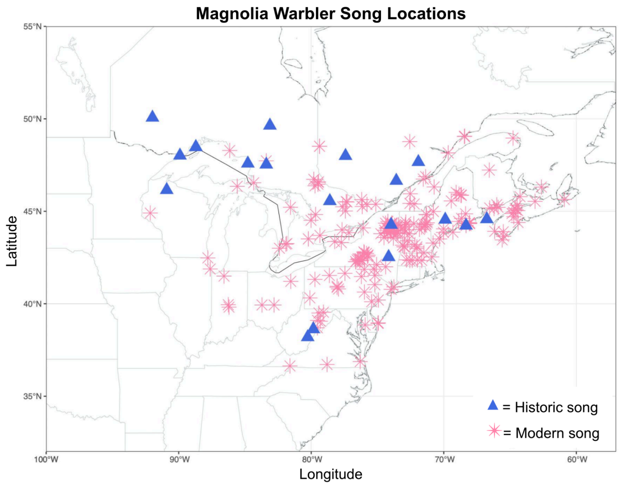
Magnolia Warbler Song Locations
55°N
50°N
45°N
40°N
35°N
Latitude
100°W
90°W
80°W
70°W
60°W
Longitude
= Historic song
= Modern song

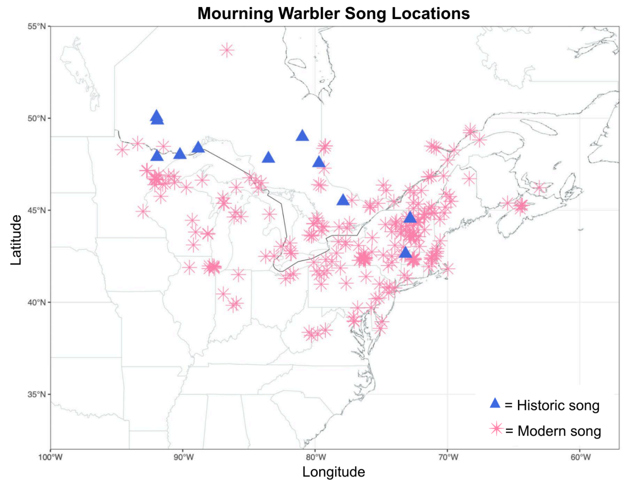
Mourning Warbler Song Locations
55°N
50°N
45°N
40°N
35°N
Latitude
100°W
90°W
80°W
70°W
60°W
Longitude
= Historic song
= Modern song

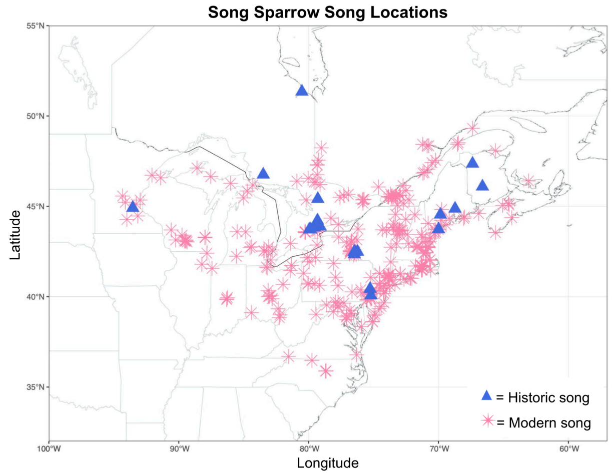
Song Sparrow Song Locations
55°N
50°N
45°N
40°N
35°N
Latitude
100°W
90°W
80°W
70°W
60°W
Longitude
= Historic song
= Modern song

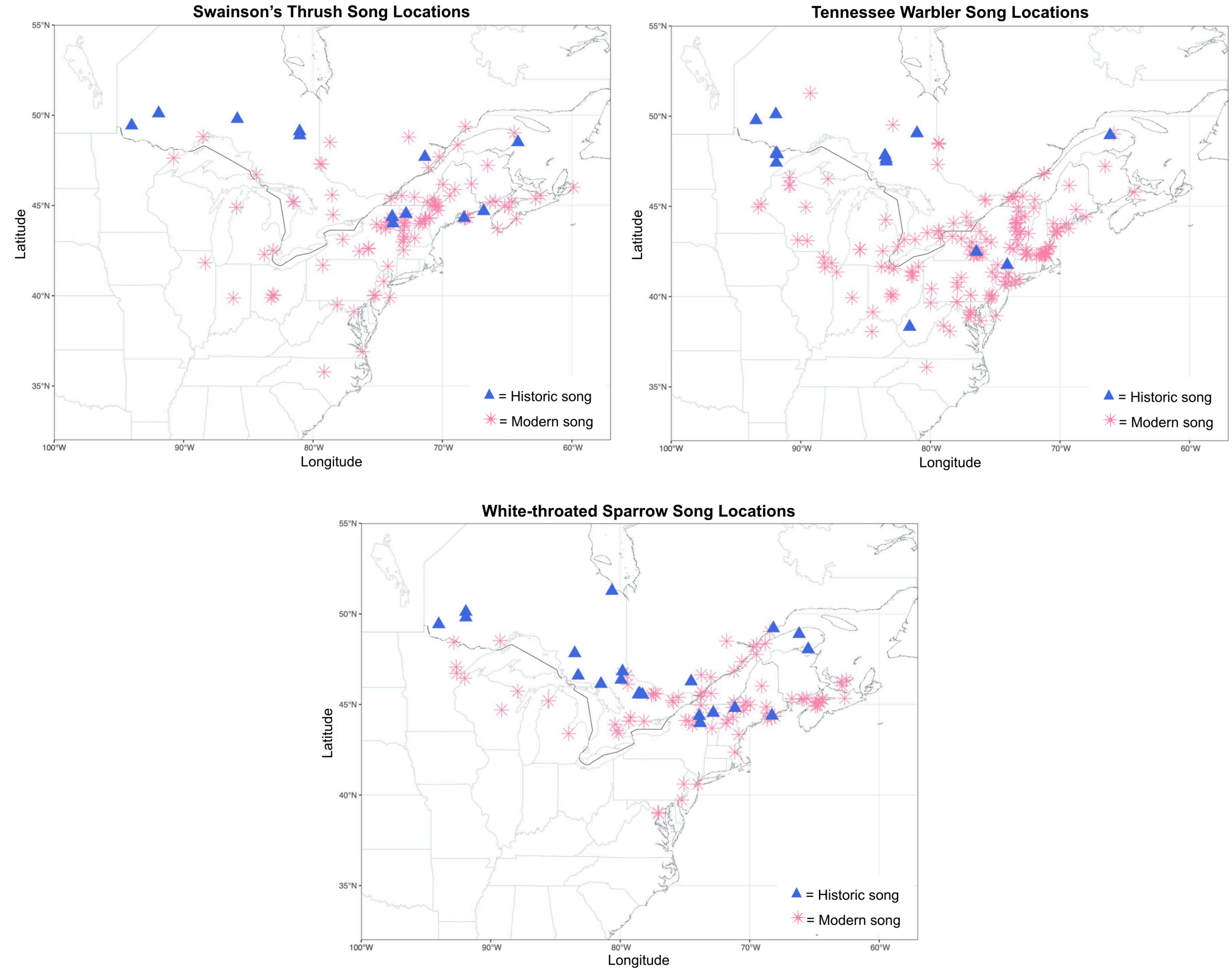


**Figure S2** Recording locations of the historic and modern songs used for each species. These maps were created with Natural Earth in R: https://www.naturalearthdata.com/about/terms-of-use/.

**Table S3** The proportion of song-feature variance that can be explained by principal component 1 (PC1) and principal component 2 (PC2) for each species.

| Species | PC1 Proportion of Variance (%) | PC2 Proportion of Variance (%) |
|---|---|---|
| Alder flycatcher | 45.25 | 20.94 |
| Chestnut-sided warbler | 34.35 | 27.20 |
| Common yellowthroat | 40.65 | 17.27 |
| Magnolia warbler | 42.72 | 19.49 |
| Mourning warbler | 41.84 | 17.81 |
| Song sparrow | 36.56 | 21.42 |
| Swainson's thrush | 37.91 | 24.62 |
| Tennessee warbler | 33.09 | 24.04 |
| White-throated sparrow | 45.24 | 21.12 |

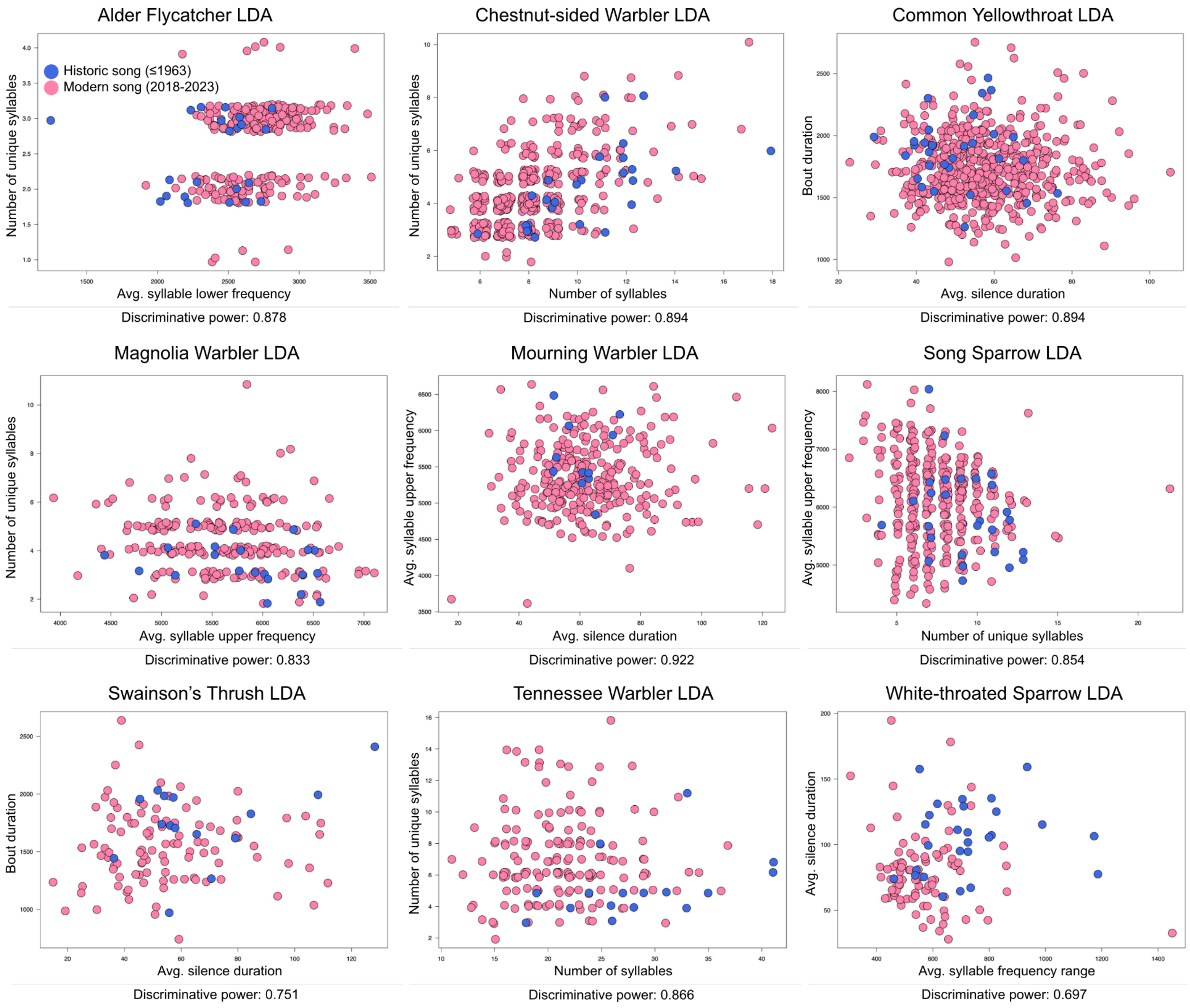


**Figure S3** Linear discriminant analysis (LDA) results for each species. The song features best able to separate the data are displayed on the graph axes and vary between species. The discriminative power of each pair of song features is listed below the graphs.

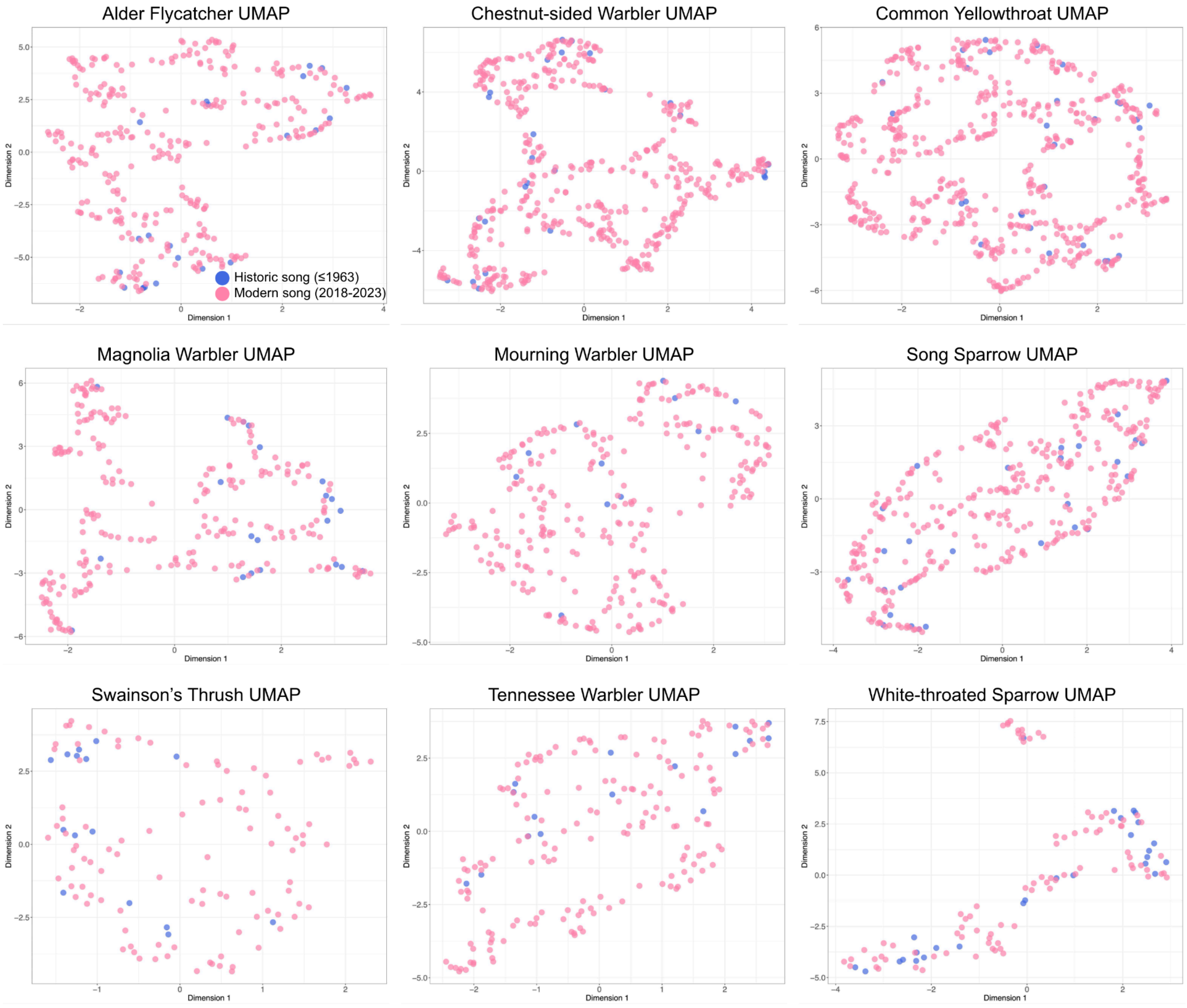


**Figure S4** Uniform manifold approximation and projection (UMAP) results for each species. Song-feature data of historic and modern groups do not cluster in dimensional space.

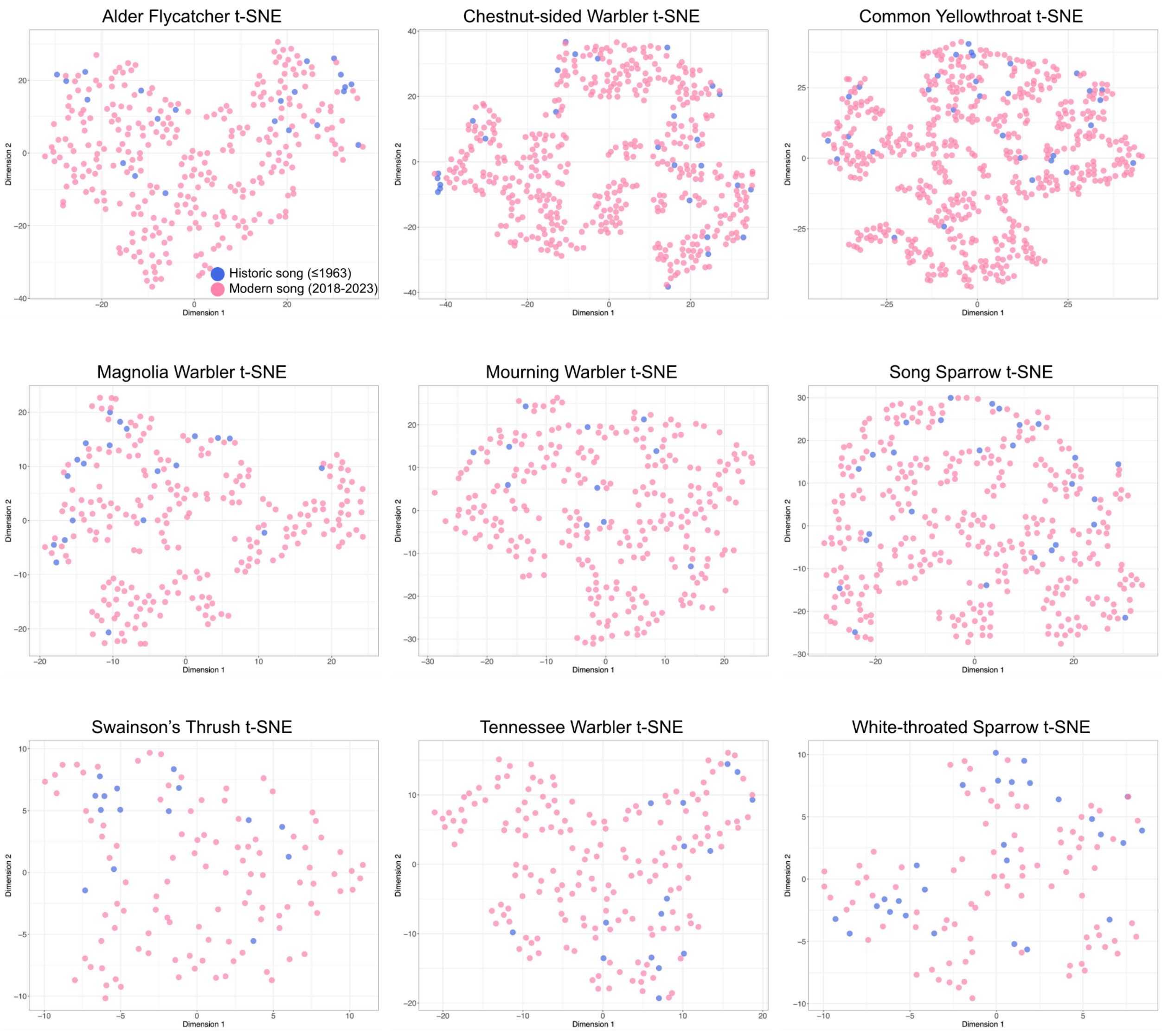


**Figure S5** t-distributed stochastic neighbor embedding (t-SNE) results for each species. Song-feature data of historic and modern groups do not cluster in dimensional space.

**Table S4** Details of the significant effects that change in population density and change in nighttime light intensity had on modern song features, with historic song features as a reference group. These were found by fitting generalized linear models to species' song features using timepoint (historic or modern), change in nighttime light intensity, and change in population density as main effects. Interaction terms were included between timepoint and each of the urbanization metrics.

| **Species** | **Song Feature** | **Urbanization metric** | **GLM *P*-value** | **GLM *t*-value** | **Interpretation** |
|---|---|---|---|---|---|
| Alder flycatcher | Avg. syllable duration | Change in population density | $7.93 \times 10^{-5}$ | -3.998 | Increasing population density is associated with shorter average syllable duration in modern compared to historic songs. |
| Alder flycatcher | Rate of syllable production | Change in population density | 0.00587 | 2.774 | Increasing population density is associated with a faster rate of syllable production in modern compared to historic songs. |
| Magnolia warbler | Avg. silence duration | Change in population density | 0.0156 | -2.435 | Increasing population density is associated with shorter average silence duration in modern compared to historic songs. |
| Magnolia warbler | Avg. syllable lower frequency | Change in population density | 0.0441 | 2.0234 | Increasing population density is associated with greater syllable lower frequency in modern compared to historic songs. |
| Magnolia warbler | Rate of syllable production | Change in population density | 0.0228 | 2.292 | Increasing population density is associated with a faster rate of syllable production in modern compared to historic songs. |
| Magnolia warbler | Number of syllables | Change in population density | 0.0266 | 2.230 | Increasing population density is associated with a greater number of syllables in modern compared to historic songs. |
| Song sparrow | Rate of syllable production | Change in population density | 0.0249 | 2.252 | Increasing population density is associated with a faster rate of syllable production in modern compared to historic songs. |
| Song sparrow | Number of syllables | Change in population density | 0.0203 | 2.331 | Increasing population density is associated with a greater number of syllables in modern compared to historic songs. |
| Song sparrow | Number of syllables | Change in nighttime light intensity | 0.00961 | -2.604 | Increasing nighttime light intensity is associated with lower number of syllables in modern compared to historic songs. |

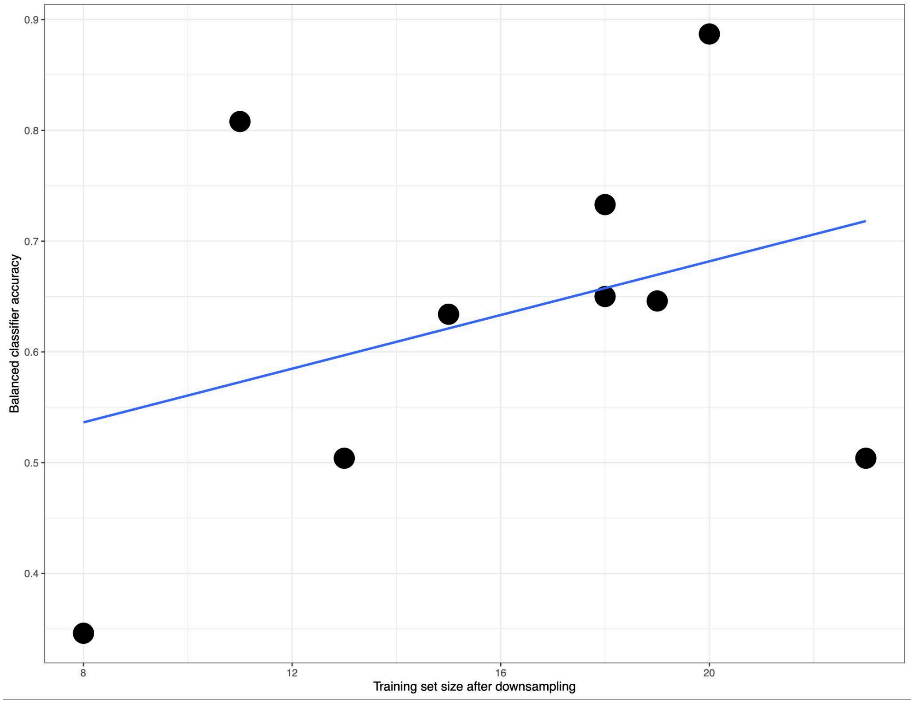


**Figure S6** The random forest classifiers' balanced accuracies are not significantly correlated with training set size after downsampling ($r$ = 0.346, $P$ = 0.361). Training set sizes varied because of the number of songs available for each species.

**Table S5** The random forest classifiers' variables of importance. Results are tallied across the nine species' random forest classifiers.

| Song feature | Most important | 2nd most important | 3rd most important | Total appearances in top 3 |
|---|---|---|---|---|
| Number of unique syllables | 0 | 3 | 2 | 5 |
| Average silence duration | 3 | 0 | 0 | 3 |
| Number of syllables | 1 | 1 | 1 | 3 |
| Average syllable lower frequency | 2 | 1 | 0 | 3 |
| Average syllable duration | 1 | 1 | 1 | 3 |
| Average syllable frequency range | 0 | 2 | 1 | 3 |
| Rate of syllable production | 1 | 0 | 2 | 3 |
| Bout duration | 1 | 1 | 1 | 3 |
| Average syllable upper frequency | 0 | 0 | 1 | 1 |

**Table S6** Correlation coefficients (*r*) between the urbanization metrics used in the GLMMs that determined whether these urbanization metrics are informative of modern-day song variation along an urbanization gradient.

| **Urbanization Metric** | Nighttime Light Intensity | Tree Coverage | Population Density | Distance to Major Road |
|---|---|---|---|---|
| Nighttime Light Intensity | | | | |
| Tree Coverage | $r$ = -0.17 | | | |
| Population Density | $r$ = 0.34 | $r$ = -0.05 | | |
| Distance to Major Road | $r$ = -0.40 | $r$ = 0.12 | $r$ = -0.12 | |
| Human Footprint Index | $r$ = 0.89 | $r$ = -0.20 | $r$ = 0.29 | $r$ = -0.46 |

**Table S7** Details of the significant GLMM models that fit modern song-feature data to distance to the nearest major road, population density, nighttime light intensity, and tree coverage.

| **Species** | **Song Feature** | **Urbanization metric** | **GLMM *P*-value** | **GLMM *t*-value** | **Correlation coefficient (*r*)** | **Correlation significance (*P*)** | **Observed relationship to urbanization** |
|---|---|---|---|---|---|---|---|
| Common yellowthroat | Avg. silence duration | Distance to nearest major road | 0.00425 | 2.859 | 0.102 | **0.0173** | **Shorter silences near major roads** |
| Common yellowthroat | Avg. syllable frequency range | Distance to nearest major road | 0.0263 | -2.221 | -0.110 | **0.0103** | **Larger syllable frequency range near major roads** |
| Common yellowthroat | Rate of syllable production | Population density | 0.0366 | 2.090 | 0.0521 | 0.224 | Faster syllable production in more populated areas |
| Common yellowthroat | Number of syllables | Population density | 0.0190 | 2.346 | 0.0480 | 0.263 | More syllables in more populated areas |
| Common yellowthroat | Avg. syllable upper frequency | Tree coverage | 0.0407 | -2.047 | -0.0876 | **0.0407** | **Higher frequencies with less tree coverage** |
| Magnolia warbler | Avg. syllable upper frequency | Population density | 0.0173 | 2.380 | 0.134 | **0.0450** | **Higher frequencies in more populated areas** |
| Mourning warbler | Number of syllables | Population density | 0.0274 | 2.206 | 0.0816 | 0.183 | More syllables in more populated areas |
| Mourning warbler | Rate of syllable production | Population density | 0.00251 | 3.022 | 0.0758 | 0.216 | Faster syllable production in more populated areas |
| Mourning warbler | Rate of syllable production | Nighttime light intensity | 0.0141 | -2.455 | -0.0982 | 0.109 | Slower syllable production in areas with more light pollution |
| Tennessee warbler | Avg. silence duration | Tree coverage | 0.0360 | 2.097 | 0.143 | 0.0639 | Shorter silences with less tree coverage |
| White-throated sparrow | Avg. syllable lower frequency | Nighttime light intensity | 0.0240 | 2.257 | 0.245 | **0.0215** | **Higher frequency in areas with more light pollution** |
| White-throated sparrow | Avg. syllable upper frequency | Nighttime light intensity | 0.0152 | 2.428 | 0.247 | **0.0203** | **Higher frequency in areas with more light pollution** |
| White-throated sparrow | Avg. syllable frequency range | Distance to nearest major road | 0.0414 | 2.040 | 0.165 | 0.125 | Smaller frequency range near major roads |
